## Supplementary Material for "Profiling the physiological impact of aberrant folded-state protein filamentation in cells"

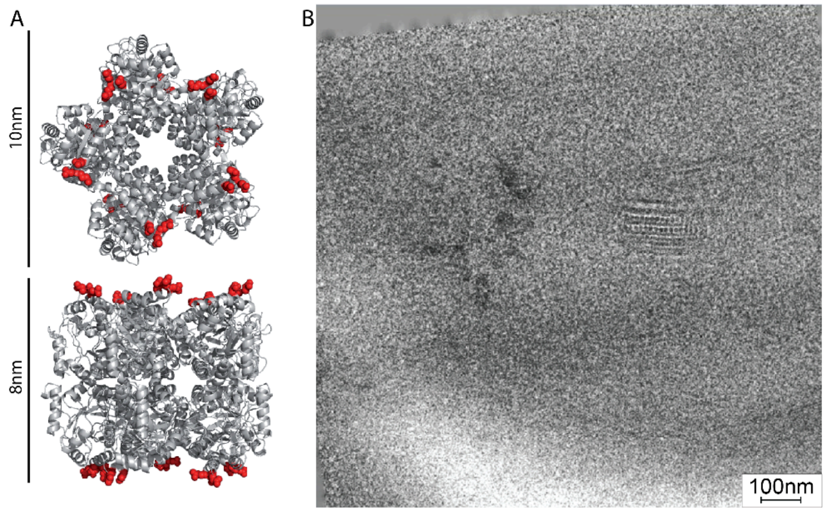


**Figure S1. Cryo-TEM of 1M3U filament shows the decamer structure. A.** The X-ray structure of 1M3U. The length and height of the decamer is indicated on the side. Mutated residues are highlighted in red. **B.** Cryo-electron Tomogram of yeast cells expressing 1M3U. Individual decamers are observed.

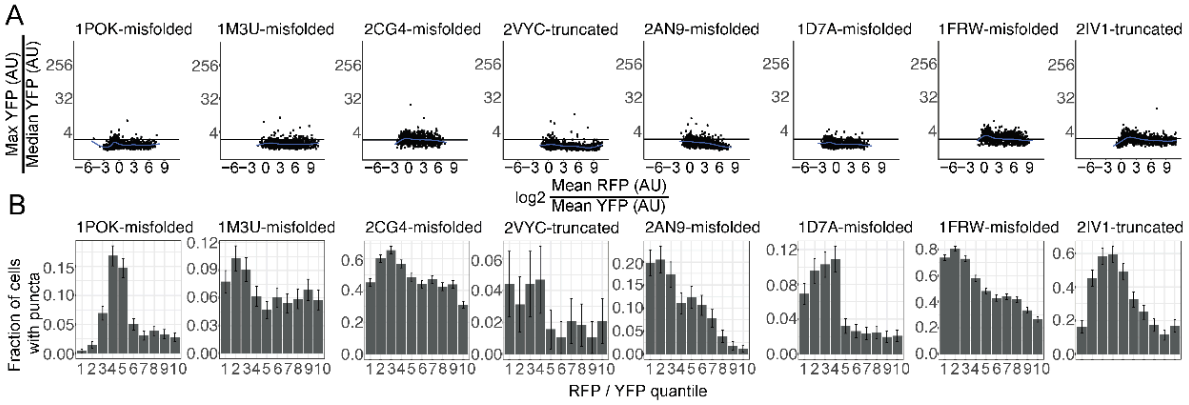


**Figure S2. Coexpression of misfolded mutants with Wild types A.** Quantification of a mutant's self-assembly depending on the relative expression of its corresponding wild type variant. Misfolded- mutants were achieved with charge mutations in the core of the protein, and truncated-, mutant (see Table S1). Mutant and wild-type relative concentration were quantified individually in each cell based on green and red fluorescence intensities. The y-axis quantifies the presence of a foci/filament in a particular cell detected by the ratio of maximal/median green fluorescence in that cell. Cells with a ratio above 2.5 (horizontal black line) are considered to contain a puncta or filament. The x-axis shows the log2 of the ratio of red/green fluorescence as a proxy of the ratio of wild-type mutant. Each dot corresponds to a cell. **B.** Same data as in panel A, where the y-axis now quantifies the fraction cells containing a foci or filament (ratio > 2.5), and the x-axis corresponds to ten quantiles of red/green intensity ratios.

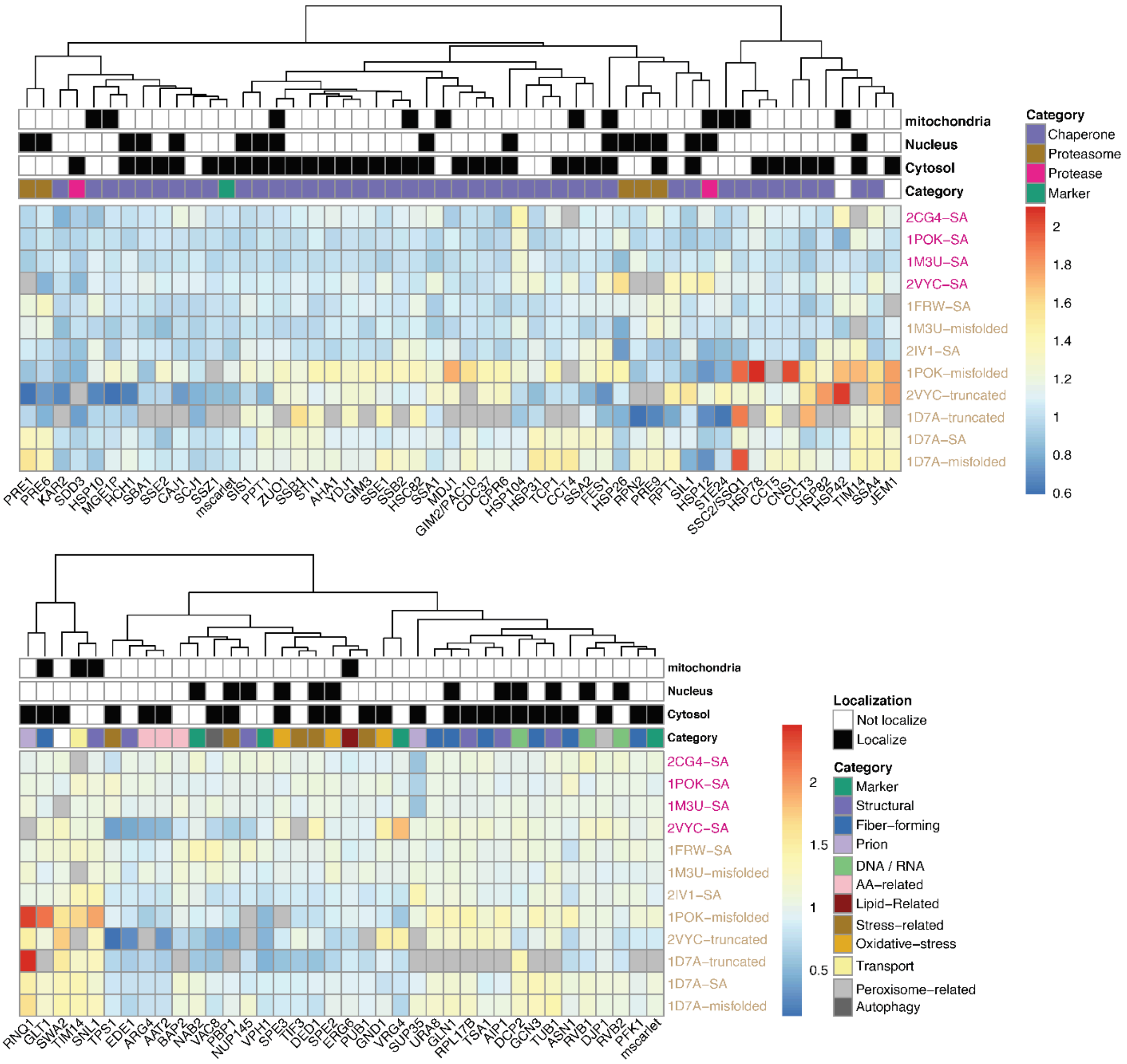

**Figure S3. Expression of yeast chaperones, proteasome, and stress-related proteins upon expression of SA and misfolded mutants.** Heatmap depicting mean Scarlet expression of cellular chaperones and proteostasis proteins (top panel) or various stress and maintenance processes (bottom panel) in cells expressing either folded (magenta) or misfolded (brown) constructs. Scarlet expression is normalized to expression of cells containing wild type variants of the same construct. Only cells detected to have a foci in the YFP channel are used. NA values appear in grey. On top localization phenotype (black) and quality control type are written.

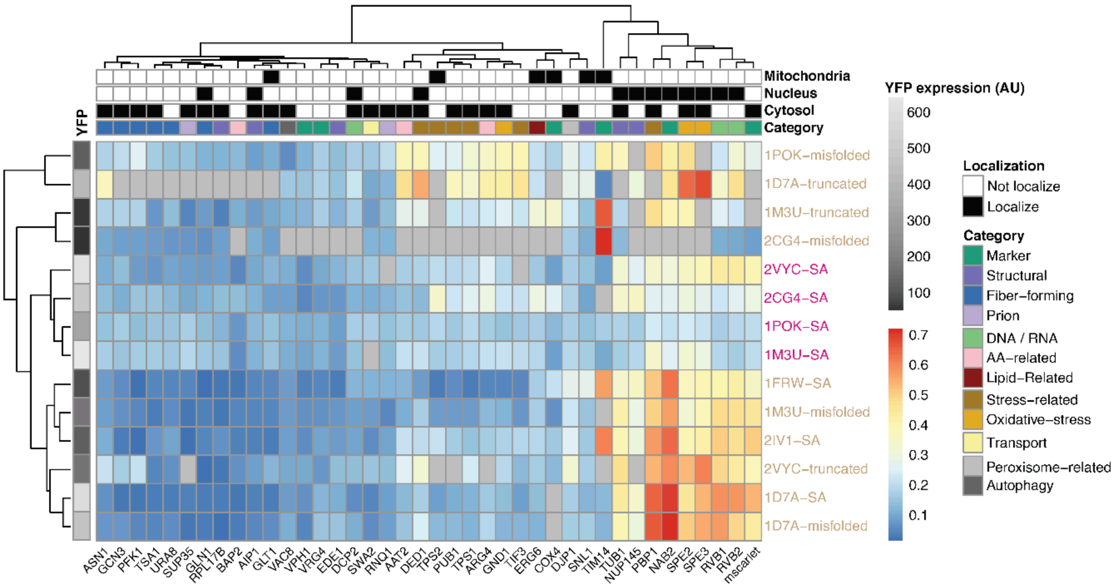


**Figure S4. colocalization 'all' categories.** Hierarchically clustered heatmap depicting colocalization scores between various stress and maintenance processes and either folded (magenta) or misfolded (brown) constructs. Only cells detected to have at least one puncta in the YFP channel are used. NA values appear in gray and reflect an insufficient number of puncta containing cells (<20) or a lack of fluorescent signal. On top localization phenotype (black) and quality control type are written. (Left), construct mean-YFP expression (AU).

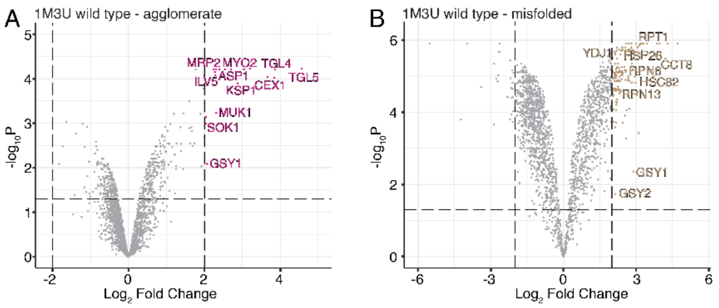


**Fig S5. Mass spectrometry following immunoprecipitation.** Mass spectrometry analysis following pull-down. **A.** Volcano plots of change in protein expression between 1M3U wild type and 1M3U agglomerate mutant. Labels are some of the significant hits. **B.** Volcano plots of change in protein expression between 1M3U wild type and 1M3U misfolded mutant. Labels are some of the significant hits.

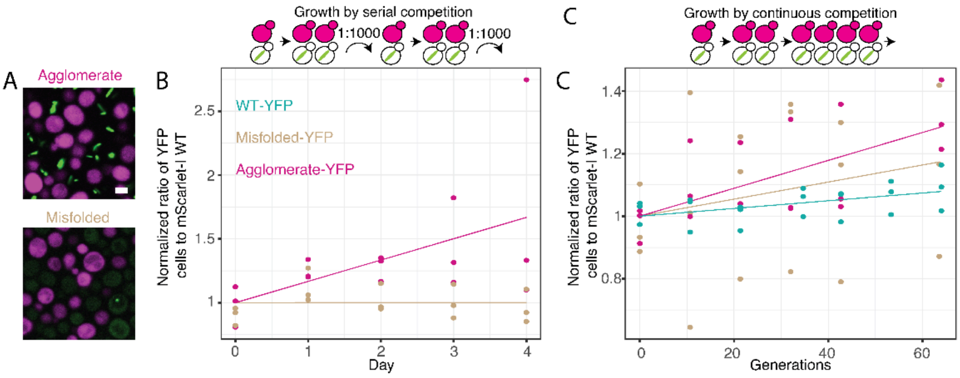


**Figure S6. Competition with WT-scarlet. A.** Representative micrographs of the competing strains. Scale bar = 3µm. **B.** Serial dilution competition assay between mat-a yeast expressing scarlet fused 1POK (magenta) and mat-a transformed with Agglomerate, or Misfolded YFP fused 1POK variants (green). Competitions occurred in liquid culture and were diluted 1000-fold daily. Every day, a sample was imaged and analyzed to quantify the populations of cells from each strain. **C.** Continuous competition that occurred over 144 hours in a chemostat.

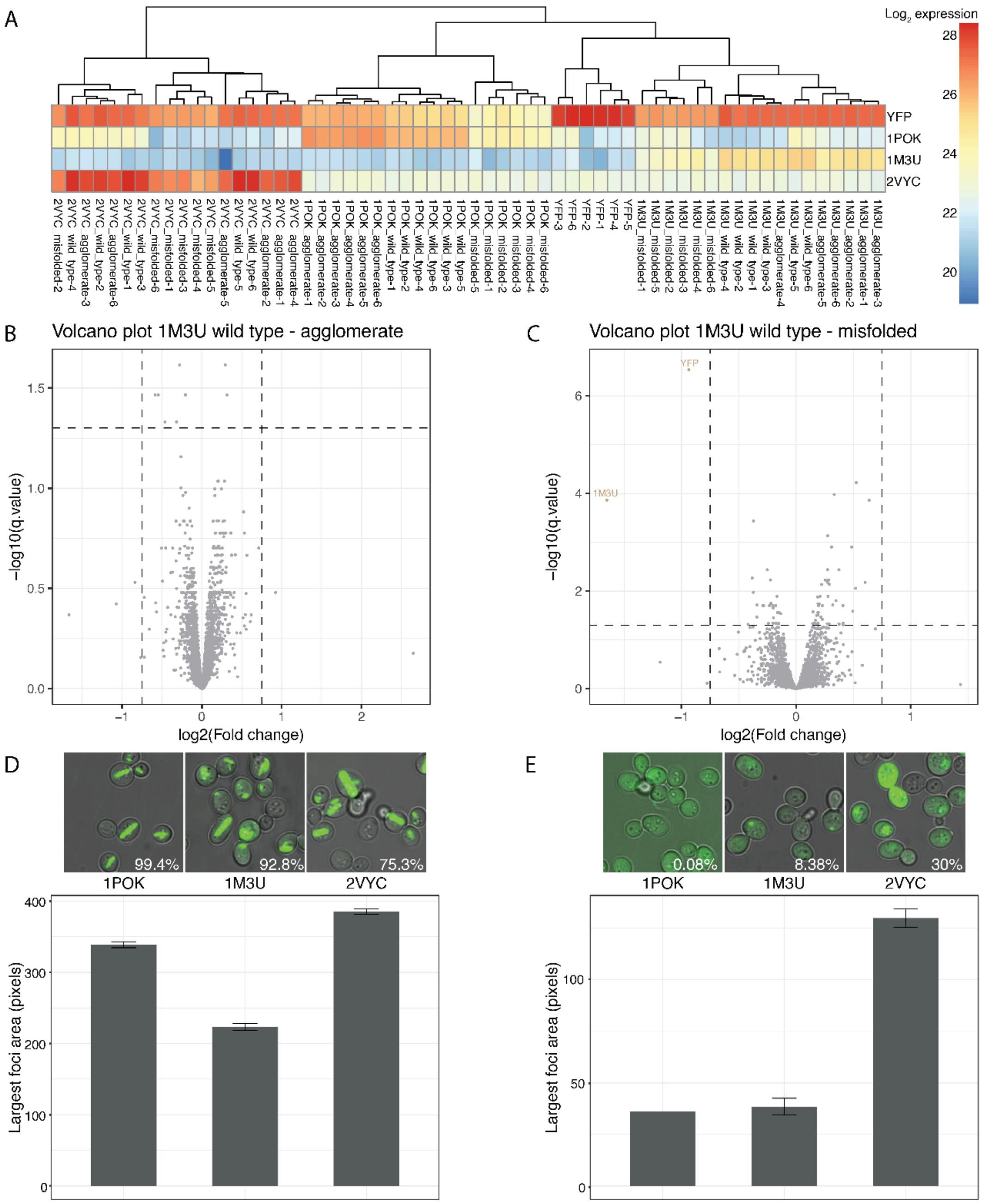


**Figure S7. Proteomics analysis A.** Heatmap of proteomics data depicting the expression of the exogenous constructs in all samples. **B-C**. Volcano plots of protein abundance differences between wild type and agglomerating (panels B) or misfolded (panel C) mutants. Each point is an average of six replicates. Pellets were frozen from a log phase liquid culture. Proteins were considered as hits if their expression changed more than 1.68 fold and had an adjusted P value < 0.05.**B.** 1M3U wild type to agglomerate. **C.** 1M3U wild type to misfolded. **D-E.** (top) representative micrographs of strain used in microscopy. Percentage of cells with foci written in white. (bottom) Quantification of the area of the largest foci in cells that contain a foci. **D.** Constructs expressing agglomerate mutants. **E.** Constructs expressing misfolded mutants.

**Table S1. Details of homomers targeted for mutations:** Details of homomers from (Garcia-Seisdedos et al. 2017). PDB code, number of subunits, symmetry, ORF, mutated positions. The linker for each protein was GGGGSGGGGS and the fused fluorophore was Venus YFP. 3N75 was removed from this study due its toxicity in yeast. 1YAC and 1L6W were removed as their filament mutants expressed poorly and cells expressing them showed pronounced autofluorescence.

| PDB code | Gene name | Protein name | Ref. | # sub units | MW | Sym | ORF | Agglomerating inducing mutations | Misfolding inducing mutations | Mutations inactivating activity | Used in this study | Reason for discarding |
|---|---|---|---|---|---|---|---|---|---|---|---|---|
| 2AN9 | gmk | Guanylate kinase | [1] | 6 | 23.6 | D3 | 2an9-Venus | D60Y/E61Y/ K63Y/E64Y | Q108K/I110K/F121K/I122K/L123K & truncation (deletion 3-181bp of 2an9 gene) | No | Yes | |
| 1D7A | purE | N5-carboxyaminoimidazole ribonucleotide mutase | [2] | 8 | 16.9 | D4 | Venus-1d7a | K11L/E22L/ E25L/D158L | A123K/G124K/1N27K/A128K & G89Stop | No | Yes | |
| 1FRW | mobA | Molybdenum cofactor guanylyltransferase | [4] | 8 | 21.6 | D4 | 1frw-Venus | D170L/D173L/ K175L/D176L | Truncation (deletion 3-271bp of 1frw gene) | No | Yes | |
| 2CG4 | asnC | Regulatory protein AsnC | [5] | 8 | 17 | D4 | Venus-2cg4 | K126Y/D131Y | C70K/I72K/G73K/I74K & K126Stop | No | Yes | |
| 1YAC | ycaC | Probable hydrolase YcaC | [6] | 10 | 22.9 | D5 | 1yac-Venus | D92Y/E94Y/ K98Y/K101Y | | No | No | Low expression of mutant |
| 1L6W | fsaA | Fructose-6-phosphate aldolase 1 | [7] | 10 | 23 | D5 | 1l6w-Venus | K97Y/K100Y/ E102Y | | No | No | Low expression of mutant |
| 2IV1 | cynS | Cyanate hydratase | Not published | 10 | 17 | D5 | 2iv1-Venus | K24L/K25L/ D26L | Truncation (deletion 3-150bp of 2iv1 gene) | No | Yes | |
| 2VYC | adiA | Biodegradative arginine decarboxylase | [8] | 10 | 84.4 | D5 | 2vyc-Venus | K491Y/D494Y/ D497Y | Truncation (deletion 3-1005bp of 2vyc gene) | T420A [9] | Yes | |
| 2WCV | fucU | L-fucose mutarotase | [10] | 10 | 15.5 | D5 | 2wcv-Venus | E77Y | Truncation (deletion 3-153bp of 2wcv gene) | No | No | Low frequency of filaments |
| 1POK | iadA | Isoaspartyl dipeptidase | [11] | 8 | 41.1 | D4 | Venus-1pok | E239Y & E239Y/ E243Y/K247Y | L354K/L356K/V357K/M358K & L240Stop | R169M [12] | Yes | |
| 1M3U | panB | 3-methyl-2-oxobutanoate hydroxymethyltransferase | [13] | 10 | 28.2 | D5 | 1m3u-Venus | D157L/E158L/ D161L | L180K/V183K/A188K/I191K Truncation (deletion 3-468bp of 1m3u gene) | D45A/D84A/E114A [13] | Yes | |
| 3N75 | ldcI | Lysine Decarboxylase | [14] | 10 | 81.2 | D5 | 3n75-Venus | D460L | | No | No | Toxic for the cell |

**Table S2 Plasmids used in this study.**

**Table S3. Proteomics raw data following bpca imputation.**

**Table S4. Manual quantifications of agglomerates in dividing cells.**

| Protein | n | % |
|---|---|---|
| Filaments passing from mother to daughter | | |
| 1pok | 83 | 30.1 |
| 1m3u | 69 | 20.2 |
| 2vyc | 59 | 20.3 |
| Filament aligned with tubulin | | |
| 1pok | 84 | 59.5 |
| 1m3u | 56 | 46.4 |
| 2vyc | 48 | 54.1 |

**Table S5. Mass Spectrometry protein hits.**

**Table S6. mScarlet-I C-SWAT mini library.**


1. Hible G, Renault L, Schaeffer F, Christova P, Zoe Radulescu A, Evrin C, et al. Calorimetric and crystallographic analysis of the oligomeric structure of Escherichia coli GMP kinase. J Mol Biol. 2005;352: 1044–1059.

2. Mathews II, Kappock TJ, Stubbe J, Ealick SE. Crystal structure of Escherichia coli PurE, an unusual mutase in the purine biosynthetic pathway. Structure. 1999;7: 1395–1406.

3. Hoskins AA, Morar M, Kappock TJ, Mathews II, Zaugg JB, Barder TE, et al. N5-CAIR mutase: role of a CO2 binding site and substrate movement in catalysis. Biochemistry. 2007;46: 2842–2855.

4. Lake MW, Temple CA, Rajagopalan KV, Schindelin H. The crystal structure of the Escherichia coli MobA protein provides insight into molybdopterin guanine dinucleotide biosynthesis. J Biol Chem. 2000;275: 40211–40217.

5. Thaw P, Sedelnikova SE, Muranova T, Wiese S, Ayora S, Alonso JC, et al. Structural insight into gene transcriptional regulation and effector binding by the Lrp/AsnC family. Nucleic Acids Res. 2006;34: 1439–1449.

6. Colovos C, Cascio D, Yeates TO. The 1.8 A crystal structure of the ycaC gene product from Escherichia coli reveals an octameric hydrolase of unknown specificity. Structure. 1998;6: 1329–1337.

7. Thorell S, Schürmann M, Sprenger GA, Schneider G. Crystal structure of decameric fructose-6-phosphate aldolase from Escherichia coli reveals inter-subunit helix swapping as a structural basis for assembly differences in the transaldolase family. J Mol Biol. 2002;319: 161–171.

8. Andréll J, Hicks MG, Palmer T, Carpenter EP, Iwata S, Maher MJ. Crystal structure of the acid-induced arginine decarboxylase from Escherichia coli: reversible decamer assembly controls enzyme activity. Biochemistry. 2009;48: 3915–3927.

9. Hong EY, Lee S-G, Yun H, Kim B-G. Improving the stability and activity of arginine decarboxylase at alkaline pH for the production of agmatine. Front Catal. 2021;1. doi:10.3389/fctls.2021.774512

10. Lee K-H, Ryu K-S, Kim M-S, Suh H-Y, Ku B, Song Y-L, et al. Crystal structures and enzyme mechanisms of a dual fucose mutarotase/ribose pyranase. J Mol Biol. 2009;391: 178–191.

11. Jozic D, Kaiser JT, Huber R, Bode W, Maskos K. X-ray structure of isoaspartyl dipeptidase from E.coli: a dinuclear zinc peptidase evolved from amidohydrolases. J Mol Biol. 2003;332: 243–256.

12. Kime L, Vincent HA, Gendoo DMA, Jourdan SS, Fishwick CWG, Callaghan AJ, et al. The first small-molecule inhibitors of members of the ribonuclease E family. Sci Rep. 2015;5: 8028.

13. von Delft F, Inoue T, Saldanha SA, Ottenhof HH, Schmitzberger F, Birch LM, et al. Structure of E. coli ketopantoate hydroxymethyl transferase complexed with ketopantoate and Mg2+, solved by locating 160 selenomethionine sites. Structure. 2003;11: 985–996.

14. Kanjee U, Gutsche I, Alexopoulos E, Zhao B, El Bakkouri M, Thibault G, et al. Linkage between the bacterial acid stress and stringent responses: the structure of the inducible lysine decarboxylase. EMBO J. 2011;30: 931–944.
